# SNaReSim: Synthetic Nanopore Read Simulator

**DOI:** 10.1101/133652

**Authors:** Philippe Faucon, Parithi Balachandran, Sharon Crook

## Abstract

Nanopores represent the first commercial technology in decades to present a significantly different technique for DNA sequencing, and one of the first technologies to propose direct RNA sequencing. Despite significant differences with previous sequencing technologies, read simulators to date make similar assumptions with respect to error profiles and their analysis. This is a great disservice to both nanopore sequencing and to computer scientists who seek to optimize their tools for the platform. Previous works have discussed the occurrence of some k-mer bias, but this discussion has been focused on homopolymers, leaving unanswered the question of whether k-mer bias exists over general k-mers, how it occurs, and what can be done to reduce the effects. In this work, we demonstrate that current read simulators fail to accurately represent k-mer error distributions, We explore the sources of k-mer bias in nanopore basecalls, and we present a model for predicting k-mers that are difficult to identify. We also propose a new SNaReSim, a new state-of-the-art simulator, and demonstrate that it provides higher accuracy with respect to 6-mer accuracy biases.

## I. INTRODUCTION

DNA sequencing has become an integral component of biological research, with applications ranging from gene network or organism identification through biological engineering. DNA sequencing has historically been dominated by synthesis-based approaches involving the replication of an existing DNA or cDNA molecule, with the attachment of fluorescent probes during synthesis for visualization of the base being added in a 4-dimensional color space. These approaches tend to have a bias in the reads selected, and tend to have reduced accuracy near the beginning and end of reads, however there tends to be minimal error bias within a single read [1]. Conversely, nanopore reads escape the read selection bias due to a lack of pre-amplification, but they provide only a single dimension of measurement, resulting in a significant within-read bias [2]. This bias is unaccounted for and lacks a published model, but it has significant implications for current generation read aligners [3]–[5] which rely on perfect or nearly perfect “seed” sequences. Unfortunately, this behavior is unaccounted for in current generation read simulators.

We demonstrate that in fact there is a read bias, that is more pronounced than the previously observed homopolymer bias, and that current generation simulators are unable to properly model this distribution. We propose a simulator based on a modified Markov chain that allows for the mutation of a reference sequence in a way that significantly improves simulation fidelity. To accomplish this task, we identify the accuracy of individual k-mers and their behavior in different contexts. We also predict key features of the error bias, and calculate their individual contributions to the final error. Our simulator is fully automated, allowing for both in silico amplification of read data, or simulation of data on completely novel genomes. Our contributions are summarized below:

- We demonstrate that while some base calling error is random there is a significant component that is systematic and unaccounted for (section IV-A).
- We develop a set of features for predicting k-mer accuracy and show that our model generally correlates well with data found empirically (section IV-B).
- We propose an algorithm for simulating reads, a variant of the Hidden Markov Model employed by Nanosim [6], with modifications to apply observed k-mer bias. We demonstrate that it models the k-mer error distribution better than other popular read simulators [6]–[8] (section IV-C).

## II. RELATED WORK

Cost, throughput, and accuracy have been major hindrances in DNA sequencing. With the development of NGS technologies cost has reduced while throughput and accuracy continue to climb. Still, simulators offer a significant benefit due to their low cost and exceptionally high throughput, allowing testing while developing new algorithms. Simulators aim to produce sequences with the most fidelity possible for their given platform. As such, simulated reads should account for biological and technical bias [9].

Simulators typically generate synthetic reads by extracting a sequence from a reference genome and then introducing errors into that sequence. Parameters required by simulators to introduce these features into the sequence are either provided at run time or are stored inside a metadata called a model or error profile. By analyzing the alignment of empirical data to a reference genome error profiles are created. Errors have been generated by first predicting a quality score [10], by base position within a simulated read [11], or by predicting an error sequence and then applying it to a read [6], [7], [12].

Modeling of third generation single-molecule reads has many advantages when compared with second generation sequencers. Polymerase chain reaction(PCR), required by second generation sequencers during the pre-amplification step, introduces significant bias, but is not required for third generation sequencers [13]. GC bias, a secondary effect of the PCR amplification step, resulting in low base accuracy and high coverage variability, is also removed [14]. Third generation sequencing on the other hand has new challenges that must be modeled. Simulators for third generation sequencers must deal with longer reads containing significantly larger stretches of errors, and a biased error that varies with nearby nucleotides. There are two main platforms for third generation sequencing, PacBio's SMRT sequencing, and Oxford Nanopore's nanopore sequencing. Each platform has their own read simulators, but none are unable to properly model the k-mer bias of nanopore sequencing data.

NanoSim [6] is a read simulator for ONT data, modeling reads as the result of a Hidden Markov model. The model is fit by aligning empirical reads to a reference genome, then collecting a list of error subtypes, lengths, and transition probabilities. Reads can be simulated by sampling a length, then generating a sequence of errors. One major limitation of this approach is that it is unable to properly model k-mer bias; this is somewhat overcome by a post-processing step where all homopolymers of length greater than 5 (e.g. “AAAAAAA”) are compressed to a length of 5.

LongISLND [8] is a read simulator developed for PacBio data. It models k-mer bias explicitly and directly by storing observed mutations for each k-mer it sees, stored as a key-value pair. It also uses an Extended K-mer (EKmer) model to aid in homopolymer error generation. An EKmer is a regular k-mer followed by an integer representing a length of homopolymer covering the middle term in the k-mer facilitating the simulation of arbitrary stretches of homopolymer without explicitly observing them. One major limitation of LongISLND is that while is is able to approximate the k-mer bias it lacks the accuracy level observed in true ONT data. This is likely due to ONT data having grouped errors (i.e. the probability of error given an error exists nearby is higher than normal) that are not properly modeled using their simulator.

## III. PROBLEM DESCRIPTION

DNA has a label space of Σ ∈ {*A, C, T, G*} representing the different nucleotide bases. Let *s* ∈ Σ*^n^* be a DNA strand of length *n*. Using Nanopore technology, a series of discrete measurements *δ* is generated, representing the current that passes through the nanopore at each time step from time *t*_0_ to *t_T_*, where *t_T_* reflects the total time required for *s* to completely transit the Nanopore. This creates a corresponding vector Δ ∈ *R^d^*, where *d* >> *n* is the number of discrete measurements obtained from time *t*_0_ to *t_T_*.

The vector Δ is subsequently binned into *q* different bins, *B* = *b*_1_, *b*_2_…*b_q_*, using different time intervals that capture ≈ 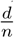 measurements per bin. The mean *µ_i_* and standard deviation *σ_i_* are subsequently derived for each bin *b_i_*. An approximation strand, 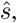 of the original strand *s* is reconstructed or “base called” using the sequence of (*µ_i_*, *sigma_i_*) pairs using either a Recurrent Neural Network [15], or a Hidden Markov Model [16], [17] in conjunction with a lookup table from the Nanopore manufacturer that specifies the most probable k-mer base pair for the given *µ_i_* value. Using the R9 pore from Oxford Nanopore *µ_i_* is best explained by a sequence of 6 DNA bases, thus the k-mer length *k* = 6 is used in later analysis.

### A. Error Source Identification

Naively, one may assume that error is distributed evenly over 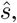, however it is trivial to see that it is not the case. The expected mean current *µ* exists on a linear range, thus k-mers near the minimum current are not as influenced by *δ* readings lower than their expected mean, with the opposite true for readings near the top of the range. Second, if the density range of the k-mer currents is uneven, more error is to be expected in high density regions [2]. Third, some sequences of K-mers are easily identifiable due to large changes in current, while others are difficult to identify due to small changes [2].

To elucidate the drivers of k-mer bias we first calculate a list of k-mer accuracies *K*, and propose a set of features *F*, where each feature *f_i_* confers an increase or decrease to the accuracy of each k-mer. Error cannot be negative, thus we can approximate the influence of each feature by solving Equation 1.

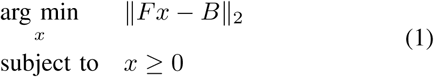

Where x is a vector indicating the contribution of each factor F. Fx is then the best approximation of the original k-mer bias with respect to euclidean distance.

### B. Read Simulation

Read simulation occurs in 2 phases: model fitting, then simulating reads from an input genome. Fitting the model requires existing reads to be aligned to a reference genome, alignment is performed using BWA-MEM [4] with standard options. From each read, the top alignment is selected, and unaligned reads are discarded. From the remaining reads, parameters are extracted with respect to average error length for the error types {Insertion, Deletion, Mismatch}, transition probabilities, and the accuracy of all k-mers of length 6. The inverse of the k-mer accuracy is also used as a “cost” of correctly predicting a k-mer. These parameters are then stored for downstream simulation.

Reads are simulated according to a hidden-markov model (HMM) which generates transitions between correct and erroneous stretches, and the lengths of those stretches. This approach has the benefit of easy explainability, but has limitations in that it is unable to correctly model k-mer accuracy distribution. Moreover, this shortfall is difficult to overcome, as modeling k-mer accuracies as states would require an intractable number of parameters. To remedy this situation we allow errors and error lengths to be generated according to the HMM, but we adjust the lengths by using a cost function described above in a process detailed in Algorithm 1.

**Algorithm 1:**
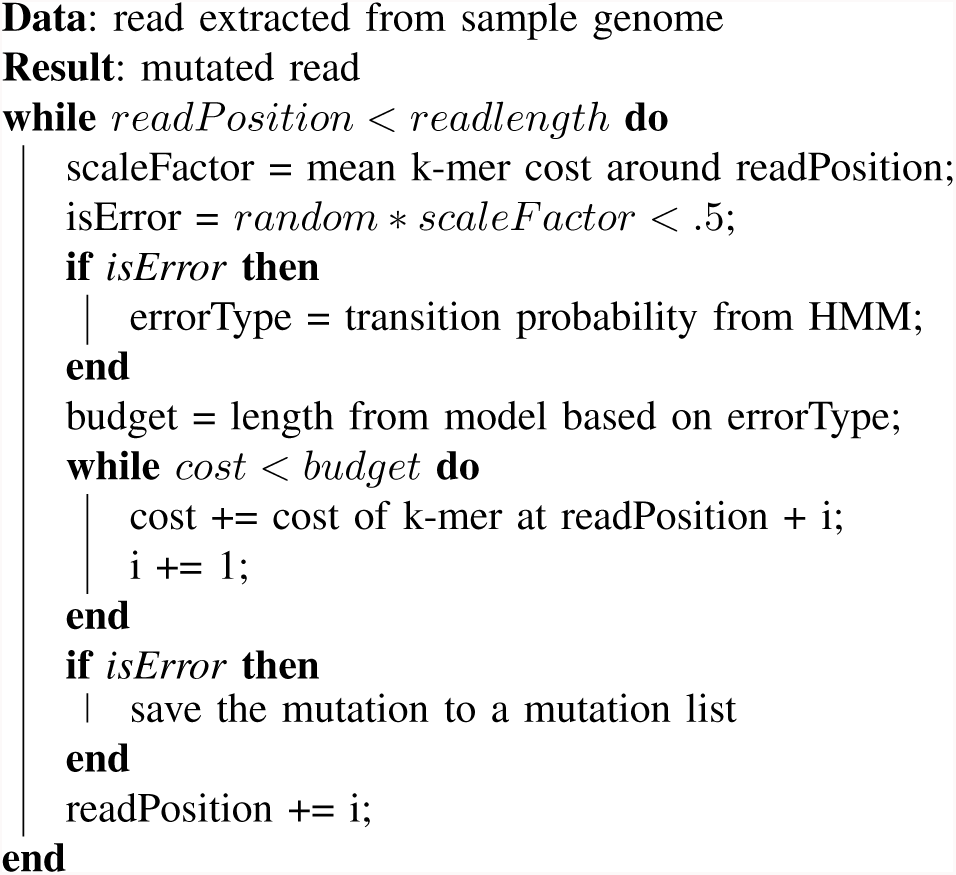
Generation of mutation list for sampled read using the k-mer biased HMM model.

## IV. RESULTS

### A. Modeling K-mer Bias

Before attempting to model k-mer bias in real data is it important to understand the granularity at which it occurs, and the consistency. To this end we examined 2 public datasets ^1^ to determine whether bias was present, and to what extent. We show that while there is a consistent k-mer bias within experiments, there are significant differences between the R7 and R9 pores in Figure 1. We attribute the majority of the differences to the pores themselves, but some influence also likely comes from different versions of ONT basecaller used on each dataset.

**Fig. 1:**
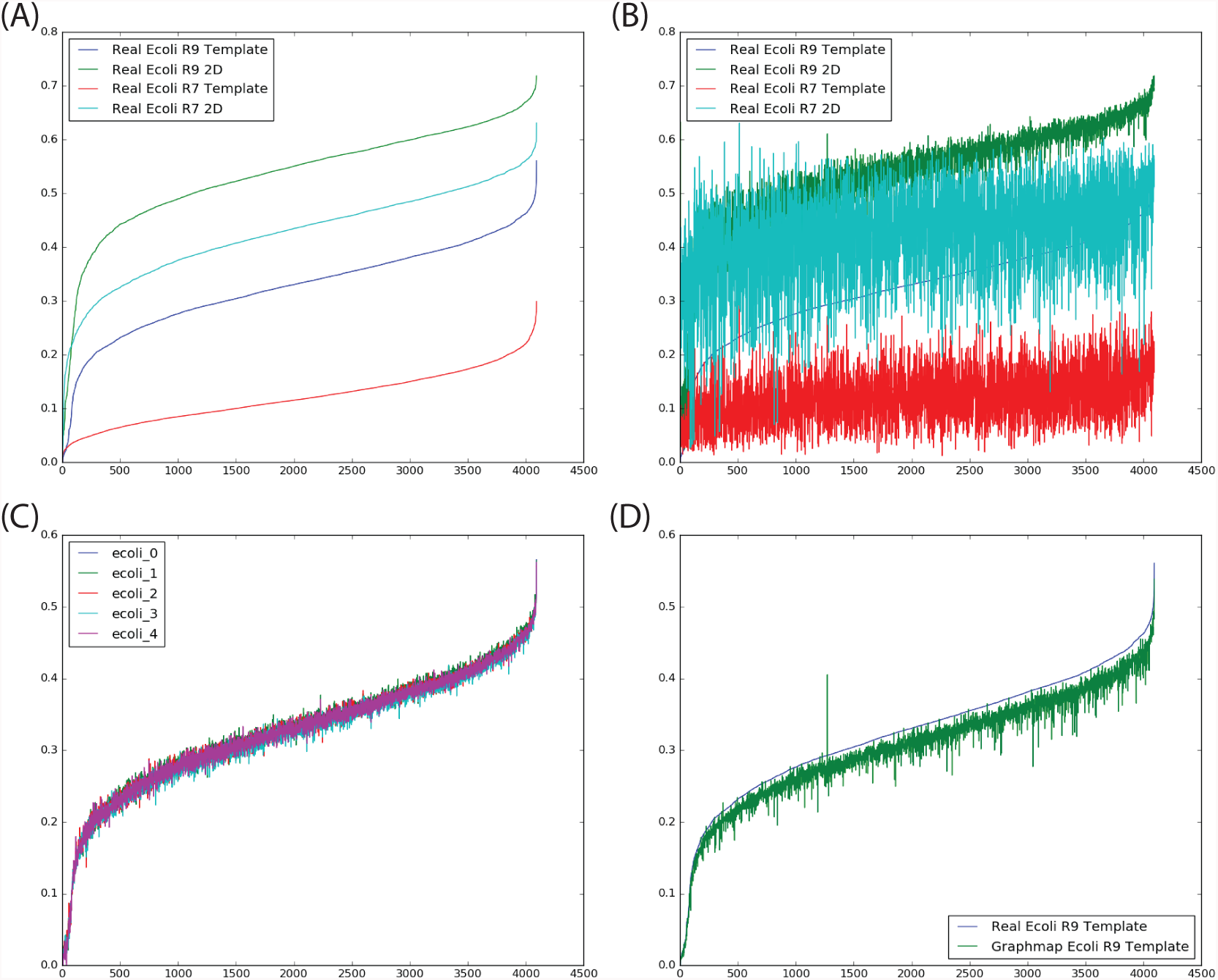
The accuracies of all 6-mers with: (A) Experiments all 6-mers sorted by accuracy with reference to themselves (B) Experiments sorted with respect to Ecoli R9 2D (C) A single experiment divided randomly into 5 groups, sorted with respect to the first group (D) only R9 ecoli template results when comparing between aligners

### B. Identifying Bias Sources

After identifying the presence of k-mer bias we attempted to elucidate the source of the bias. To this end we generated a set of features with each providing a score for each k-mer, and we attempted to find a linear combination of each feature capable of explaining the bias, and providing a fractional contribution of each step. As shown in Table I multiple error sources influence the accuracy fraction of each k-mer. We find that the strongest feature for predicting the accuracy is the median accuracy of neighbors at one step away. While this is not directly helpful, it does suggest that direct modeling of error sources must incorporate neighbor accuracy aspects. Overall we find that our predictive model provides a good first attempt, providing some insight, but leaving much of the error signal uncaptured, as illustrated in Figure 2. We expect that much of the remaining error signal is a result of missing features that may be identified through a more thorough analysis. We also expect that some of the error signal cannot be captured through linear combinations. For example, the original basecallers were incapable of capturing homopolymers with a length greater than 5, using linear combinations the k-mer “AAAAAA” would need to have 0 accuracy across all features, an unrealistic expectation.

**Fig. 2:**
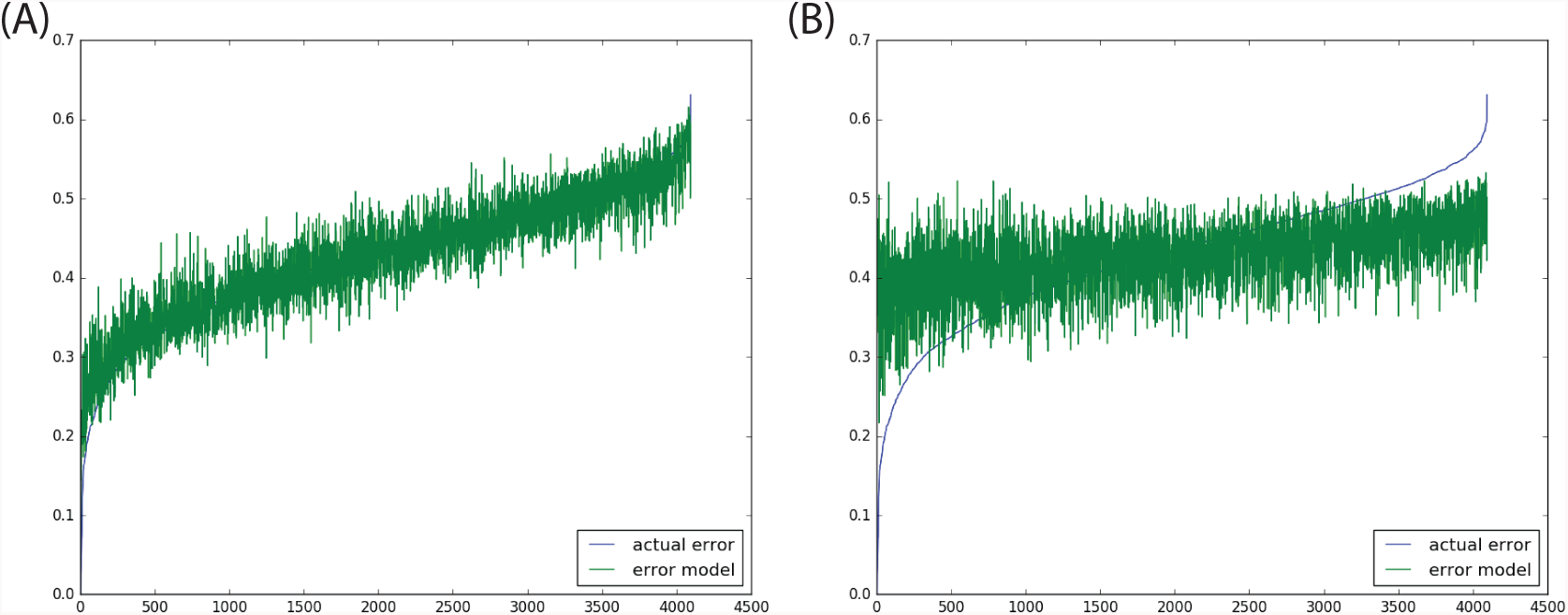
The accuracy of k-mer fit to the actual distribution from the R9 pore model. (A) Including the true accuracy of neighboring k-mers as a feature (B) using only the features in Table I

**TABLE I:** Identified error sources and their contribution to k-mer overall accuracy with respect to Figure 2

| Bias Source | Example Low | Example High | Contribution |
| --- | --- | --- | --- |
| Transition Identifiability | AAAAAA | ATATAT | 64.6% |
| Sequence Identifiability | CTGTCA | TATATT | 3.32% |
| Standard Deviation | CTAGAG | TTGAAA | 1.79% |
| Fraction A | CGTCGT | AAAAAA | 9.73% |
| Fraction T | CGACGA | TTTTTT | 2.90% |
| Fraction C | AGTAGT | CCCCCC | 9.33% |
| Fraction G | CATCAT | GGGGGG | 7.03% |

### C. Simulation Results

To validate our simulation results we generated read profiles for Nanosim [6], PBSim [7], LONGIslnd [8], and the two simulators we have proposed. Nanopore sequencing experiments typically yield between 10,000 and 20,000 reads [6], with measured statistics being approximately identical at even 20% of this size as shown in Figure 1(C). To measure this effect in simulations, we generated 2 data sets with each simulator: one with 4,000 reads, and one with 20,000 reads. These reads were then aligned using BWA-MEM [4] with standard parameters. The sum of squares error is the sum of the difference between the model 6-mer accuracy and the simulated k-mer accuracy, for all 6-mers. Results of both large and small simulations are shown in Table II, while the 20,000 read simulations are shown in Figure 3. Ultimately we find that sample size has very little influence on existing simulators, as the k-mer bias is either completely un-modeled, or has systematic difficulties capturing the k-mer error rate (LongISLND). In our simulators we find some improvement through increased sample size, though most of the remaining difference does appear to be from systematic errors.

**Fig. 3:**
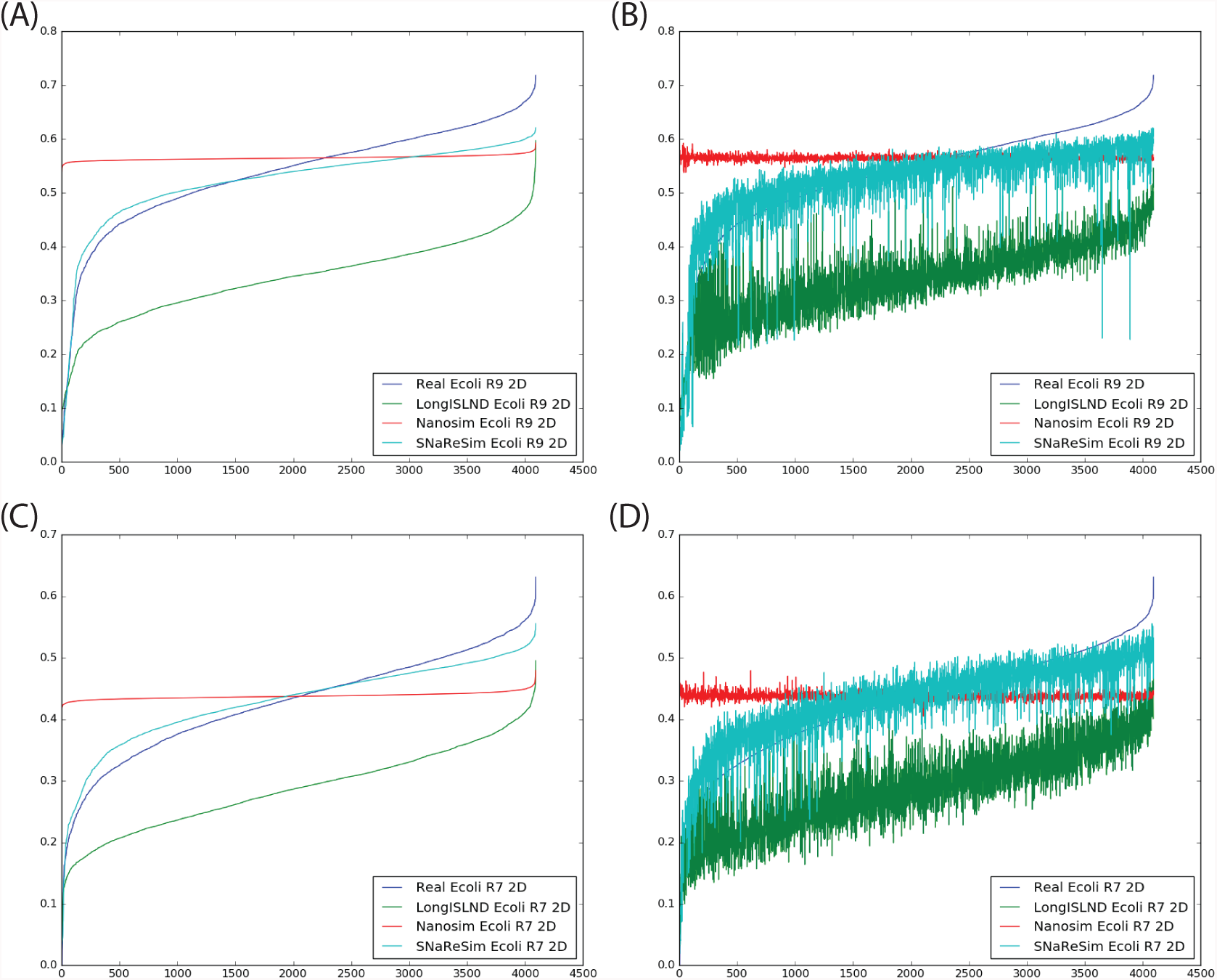
Comparison of 3 previously published sequencing simulators with the new model we proposed. Fit of R9 pore model k-mer accuracies sorted internally(A), and by 2D Ecoli, the training model(B). Fit of R7 pore model k-mer accuracies sorted internally(C), and by 2D ecoli, the training model (D).

**TABLE II:** Sum of squares error(SSE) between k-mer simulators and their training, and between the fragmented R9 data set and the complete one(Figure 1)

| Source | Sample Size | SSE |
| --- | --- | --- |
| Split R9 | 6,481 | 0.121 |
| LongISLND R9 | 20,000 | 167.04 |
| LongISLND R9 | 4,000 | 167.80 |
| Nanosim R9 | 20,000 | 44.26 |
| Nanosim R9 | 4,000 | 46.76 |
| SNRG R9 | 20,000 | 9.52 |
| SNRG R9 | 4,000 | 9.66 |
| LongISLND R7 | 20,000 | 88.78 |
| LongISLND R7 | 4,000 | 88.96 |
| Nanosim R7 | 20,000 | 30.76 |
| Nanosim R7 | 4,000 | 36.16 |
| SNRG R7 | 20,000 | 5.24 |
| SNRG R7 | 4,000 | 5.48 |

## V. CONCLUSION

We have demonstrated the presence of biased k-mer accuracy within Oxford Nanopore Technologies sequencing platform. We show that this bias is consistent within experiments, and to a large extent between sequence aligners, but varies between pore models. This information is of significant value to sequence aligners, many of which look for perfectly matching seed sequences. Due to the bias in k-mer accuracy some seed sequences are virtually impossible to find correctly, and as such should be excluded from being chosen as seeds.

We demonstrate that this k-mer bias has some observable and predictable foundation in the mean current readings of each k-mer(i.e. the manufacturers pore model). This provides a way to estimate the read accuracy for a provided pore model without performing large sequencing runs; it also suggests that a model-guided approach for nanopore design could improve overall accuracy. Alternatively it could allow for the development of specific pores, where accurate discrimination of some k-mer subtypes is more important than others.

Finally we propose a novel nanopore read simulator capable of modeling the k-mer bias observed in this experiment. We demonstrate that our simulator has a significantly reduced sum of squares error with respect to 6-mer accuracy when compared with other simulators. This provides a more realistic benchmark for sequence aligners to compare against, specifically allowing for sequence aligners tailored for nanopore sequencing data.

The availability of high accuracy reads allows for the exploration of new applications, including; sequencing of larger organisms, organism disambiguation when sequencing a population, exact sequence detection in diploid and polyploid organisms, and the ability to scaffold genomes across exceptionally long repeat regions. Providing an understanding of accuracy in nanopore design, and development of tools to aid the alignment of the produced reads is thus critical to continued progress.

## Footnotes

1 {http://lab.loman.net/2016/07/30/nanopore-r9-data-release/,http://lab.loman.net/2014/10/01/where-can-i-get-oxford-nanopore-miniontm-data-from/}

